# A haplotype-complete chromosome-level assembly of octoploid *Urochloa humidicola* cv. Tully reveals multiple genomic compositions and evolutionary histories in the species

**DOI:** 10.1101/2025.10.23.684187

**Authors:** Jose J. De Vega, Alex Durrant, Naomi Irish, Tom Barker, Rosa N. Jauregui

## Abstract

We developed a haplotype-resolved, chromosome-scale genome assembly of the *Urochloa humidicola* (Rendle) Morrone & Zuloaga cultivar Tully, an apomictic C4 forage grass cultivated in the tropics worldwide. We assembled a 4.1 Gb genome into 48 chromosomes (2n = 8x = 48), capturing 99.5% of the BUSCO markers, and annotating 259,254 protein-coding genes. Subgenome assignment revealed an octoploid AABBBBCC structure with three ancestral lineages (A, B, C), and an aneuploid composition of 14A, 22B, and 12C chromosomes. Comparative analyses with related *Urochloa* species identified *U. dictyoneura* and *U. arrecta* as potential progenitors of the B and C subgenomes, respectively. The likely progenitor of the A subgenome remains an unknown wild species from the Humidicola clade. Analysis of LTR-retrotransposons and gene collinearity further indicated a close relationship between A and B ancestries, and a distinct evolutionary path for C. Competitive read mapping across additional *U. humidicola* accessions supported multiple evolutionary histories in the species, with AABBBB (lacking C ancestry), being the most common. We found that previously described subpopulation structures can be explained by the presence or absence of C ancestry, and that sexual *U. humidicola* are likely to be autopolyploid from the B ancestry. The genome is available as assembly GCA_965614515.2. This assembly provides the first complete reference for *U. humidicola* and reveals a multi-ancestral origin and reticulated evolution in *U. humidicola*. It provides a foundation for studying complex polyploid evolution, regulation of apomixis and biological nitrification inhibition, and molecular breeding strategies.

## Introduction

*Urochloa humidicola* is a perennial C4 tropical forage grass of major economic and ecological importance in tropical livestock systems. Native to African savannas and now widely cultivated across Latin America, Africa, Asia, and Oceania, it is particularly valued for its persistence in low-fertility and waterlogged soils, and tolerance to grazing (Ferreira et al. 2021). Despite its agronomic success, *U. humidicola* poses exceptional challenges for genetic improvement. The species exhibits complex polyploidy (Higgins et al. 2022; Tomaszewska et al. 2023), with cytotypes ranging from hexaploid to nonaploid (2n = 6x–9x = 36-54).

*U. humidicola* reproduces predominantly through gametophytic apomixis, producing clonal seeds without fertilisation (da Costa Lima Moraes et al. 2023). Apomixis has long been regarded as a transformative trait that could fix heterosis through clonal seed production, enabling elite hybrid genotypes in important seed crops to be perpetually reproduced through biotechnological approaches (Underwood and Mercier 2022). *U. humidicola* and related *Urochloa* species serve as an excellent model for exploring the genomic basis of apomixis in polyploid grasses (Cervantes-Díaz et al. 2024).

Furthermore, *U. humidicola* exhibits biological nitrification inhibition (BNI) activity, a unique physiological trait that suppresses soil nitrification and mitigates nitrogen losses following fertilising (Nuñez et al. 2018; Egenolf et al. 2020; Vázquez et al. 2020). Understanding the genetic basis of BNI offers a promising route to develop biological inhibitors to reduce nitrogen leakage in arable systems (Issifu et al. 2024).

*U. humidicola* holds significant potential for climate-smart livestock production. Molecular breeding, such as marker-assisted and genomic selection, can accelerate the development of cultivars with improved nutritional quality and increased productivity (Ferreira et al. 2021). The intensification of livestock systems can reduce the land footprint of pasture expansion in the tropics, and enhanced fibre digestibility in *Urochloa* forages has been linked to lower methane emissions from ruminants, aligning breeding goals for this species with global targets to decrease greenhouse gas emissions from agriculture (Arango et al. 2020).

Here, we report a haplotype-complete, chromosome-level genome assembly of the apomictic *U. humidicola* cv. Tully. This reference genome provides the first comprehensive view of the subgenomic structure and evolutionary relationships within this complex polyploid species. It represents a valuable resource for the forage genomics community, enabling future studies on genome evolution, the genetic control of apomixis and BNI, and the deployment of genomic tools for breeding improved climate- resilient forages.

## Methods & Materials

### Sample collection, DNA extraction and sequencing

*U. humidicola* cv. Tully plants were grown for 8 weeks before DNA was extracted from seeds of accession APG 58384, obtained from the Australian Pastures Genebank. Tully is an apomictic genotype found worldwide; it was introduced into Australia in 1952 by J.F. Miles as accession CPI 16707 (AusTRCF 16707) from Rietondale Experimental Stations, Pretoria, South Africa (-25.7283, 28.236, 1,300 m elevation, 700 mm rainfall). It was later introduced to Fiji and Papua New Guinea; from there, clonal material was reintroduced from Papua New Guinea to the Tully region of North Queensland in 1973, and it was officially registered for commercial release as “Tully” in 1981 (Miles 1996). The seed of cv. Tully (APG 58384) used for sequencing comes from this reintroduced material. In America, the cultivated material is known as “pasto humidicola” or “Chetumal” (CIAT 679, PI 499378, BRA 001627, INIAP-NAPO 701), which are apomictic clones of the sequenced accession.

After 8 weeks, leaf material was harvested following two days in dark conditions. Extraction of HMW DNA was performed using the Nucleon PhytoPure kit, with a variation of the recommended protocol optimised for DNA fragment size, yield and purity. 1g snap-frozen leaf tissue was ground under liquid nitrogen for 9 minutes, with the mortar refilled with LN2 between every 90 second of grinding. The resulting powder was thoroughly mixed into Reagent 1 (with 4uL of 100mg/ml RNase A) in a 15ml tube using a bacterial spreader loop. The sample was incubated on a ThermoMixer at 37C for 30 minutes. Reagent 2 was then added and the tube was gently inverted until the contents appeared homogeneous, at which point the tube was incubated on a ThermoMixer at 65C for 10 minutes.

Following the post-lysis ice incubation, 2ml chloroform and 300uL resin was added, and the sample was mixed on a 3D platform rocker for 10 minutes at room temperature. From here, the sample was centrifuged at 1300g for 10 minutes. The upper phase was mixed with 1 volume of 25:24:1 Phenol:Chloroform:Isoamyl Alcohol, mixed on a 3D platform rocker for 10 minutes at 4C, and centrifuged (3000g) for 10 minutes. The upper phase was transferred to another 15ml Falcon tube and precipitation proceeded as recommended by the manufacturer protocol. The final elution was left open in the chemical safety cabinet for 1.5 hours to allow residual ethanol to evaporate, after which the DNA sample was left at room temperature overnight.

11.5µg of gDNA was manually sheared with the Megaruptor 3 instrument (Diagenode, P/N B06010003) according to the Megaruptor 3 operations manual, with a speed setting of 31 and input of 18ng/µl in 150µl. The sheared sample then underwent AMPure® PB bead (PacBio®, P/N 100-265- 900) purification and concentration (0.45X, eluted in 95 µL of EB) before undergoing library preparation using the SMRTbell® Express Template Prep Kit 2.0 (PacBio®, P/N 100-983-900). The HiFi library was prepared according to the HiFi protocol version 03 (PacBio®, P/N 101-853-100), and the final library was size fractionated using the SageELF® system (Sage Science®, P/N ELF0001), 0.75% cassette (Sage Science®, P/N ELD7510), with a 4.55hr program. The library was quantified by fluorescence (Invitrogen Qubit™ 3.0, P/N Q33216), and the size of the library fractions was estimated from a smear analysis performed on the FEMTO Pulse® System (Agilent, P/N M5330AA).

The loading calculations for sequencing were completed using the PacBio® SMRT®Link Binding Calculator v 11.1.0.166339. The Two largest ELF fractions proceeded to sequencing as separate libraries, and Sequencing primer 3.2 was annealed to the adapter sequence of each HiFi library. Each library was bound to the sequencing polymerase with the Sequel® II Binding Kit v3.2. Calculations for primer and polymerase binding ratios were kept at default values for the library type. Sequel® II DNA internal control 3.2 was spiked into each fraction at the standard concentration before sequencing. The sequencing chemistry used was Sequel® II Sequencing Plate 2.0 (PacBio®, P/N 101-820-200) and the Instrument Control Software v11.0.1.162970. The library was sequenced on the Sequel IIe across 3 Sequel II SMRT®cells 8M. The parameters for sequencing per SMRT cell were adaptive loading (set to 0.85 target for 2 hours), 30-hour movie, 2-hour pre-extension time, and 90pM on plate loading concentration.

### Genome assembly and quality check

Unitigs and contigs were assembled from the raw HiFi reads using HiFiasm v0.20 (Cheng et al. 2021). The quality of the assembly was assessed using several measures; contiguity was evaluated with Quast v. 5.0.2 (Gurevich et al. 2013), assembly completeness was assessed with BUSCO v5.3.2 analysis (Seppey et al. 2019) using the poales v10 database; and Merqury v1.3 (Rhie et al. 2020) was used to generate kmer completeness metrics and kmer spectra plots to confirm the assembly captured all the content from the reads.

### Scaffolding through synteny within the genome

Firstly, the contigs from each haplotype assembled by HiFiasm, haplotype 1 (hap1) and haplotype 2 (hap2), were compared separately using RagTag v2.1.0 (Alonge et al. 2022) to the “primary contigs” produced by HiFiasm. These “primary contigs” were constructed by HiFiasm with the criteria of maximising continuity, resulting in longer sequences but mixed haplotypes. For each hap1 or hap2, the PAF file internally generated by RagTag (by running Minimap 2.22 with the “-x asm5” option) was visualised as a dotplot with dotPlotly, using a minimum alignment length of 10 kb. Links in the AGP file with query or target having more than one alignment were manually filtered out.

Secondly, we compared the resulting hap1 and hap2 to each other using RagTag, as before. Then generated a new dotplot from the alignment file, and manually filtered any entries from the AGP where the query or target had more than one alignment.

Thirdly, we created a synthetic genome of *Urochloa fusca* with six chromosomes (instead of the real nine) by combining chromosomes 2 and 4, chromosomes 5 and 6, and chromosomes 1 and 7 from the available *U. fusca* genome downloaded from Phytozone. We then compared haplotype 1 and haplotype 2 from the previous step to this synthetic genome and selected the *U. humidicola* sequence that showed the longest alignment to each chromosome with the synthetic *U. fusca* genome. Finally, we compared the hap1 and hap2 resulting from the second step against the six “longest” sequences from *U. humidicola* using RagTag, following the same procedure as before; first analysing each haplotype independently, then together. In total, 225 contigs were scaffolded into 48 chromosomes.

### Gene annotation

The genome was soft-masked using the EI-Repeat pipeline (EI-CoreBioinformatics). The gene annotation was completed with Braker v3.0.8 (Gabriel et al. 2024), with default options, using as inputs the soft-masked genome, the proteins from the reference genomes of *Setaria italica, S. viridis, Sorghum bicolor, Panicum halli* and *U. fusca*, all downloaded from Phytozome v13, and all RNA-seq libraries from *U. humidicola* accessions from our previous paper (Higgins et al. 2022). The resulting annotation from Braker was used as input to Mikado v2.3.3 (Venturini et al. 2018), together with the Stringtie assemblies generated by Braker and an exon/intron junctions file generated with Portcullis v1.2.4 (Mapleson et al. 2018) from the same RNA-seq libraries. The output from Mikado was used as the final gene annotation.

### Subgenome assignment and ancestry

We used SubPhaser v20230401 (Jia et al. 2022; Zhang et al. 2024) to assign chromosomes into subgenomes (A, B, C), identify and compare LTR-RTs, and compare haplotype blocks by alignment between chromosomes. Subphaser required the reference and a list of chromosomes certain to be homoelogous (e.g. A_hap1, B_hap1, C_hap1). As we were not certain of the ancestry of some chromosomes, only the 18 chrs in hap1 were included initially. Additional homoelogous triads were entered iteratively until all haplotypes were included.

We downloaded the protein sequences of the “Angiosperms 353” (A353) markers from the “Tree of Life” project (Johnson et al. 2019). We also retrieved reads from EBI’s ENA for these markers, which were generated for 18 Urochloa species in two prior studies (Baker et al. 2022; Masters et al. 2024). We pre-processed them with Trim Galore v0.6.10 (Krueger 2015) with default options and trimmed 5 bps from each read end. Using the A353 protein sequences and the clean reads for each species, we employed Hybpiper (Johnson et al. 2016) to assemble the protein “Angiosperms 353” sequences for the 18 *Urochloa* species. In parallel, we generated the cv. Tully proteome based on the gene annotation and divided it into subsets by subgenome and haplotype (A hap1, A hap2, B hap1, etc.). A hap 3 and B hap 4 were excluded, as they were not present in all six chromosomes. We then compared these subsets from *U. humidicola* against the “Angiosperms 353” targets using Diamond v2.0.9 (Buchfink et al. 2021), and selected all queries with over 90% coverage using Seqtk v1.4.122, to extract the A353 homologs in each *U. humidicola* subgenome and haplotype. Finally, these subsets extracted from *U. humidicola*, along with the 18 sets assembled by Hybpiper (all corresponding to “Angiosperms 353” homologs), were clustered and analysed with Orthofinder v3.0 to construct a phylogenetic tree, which was visualised using the online tool itol.

### Composition in other *U. humidicola* accessions

Public DNA-seq data were downloaded from EBI’s ENA, including accessions CIAT 16888 (SRR8321580), CIAT 26155 (SRR16327314), and CIAT 26146 (13 read files from PRJNA509199), which is a sexual accession. These three datasets were pre-processed using Trim Galore v.0.6.10 (default parameters), then aligned to the assembled chromosomes with Bowtie v2.4.1 using the options “-X 500 --no-unal --no-mixed --no-discordant --no-contain --no-overlap”, ensuring only concordant paired reads were retained. The aligner’s default behaviour, where reads are aligned to their “top best” position, was applied. Since our alignment parameters only permit a read pair to align to their “top best” position, this results in a competitive mapping analysis where reads mainly map to their original subgenome for composition estimation. A coverage table was generated from the alignment files using Samtools coverage v1.51.1 (Danecek et al. 2021).

We also aligned the 32 RNA-seq libraries from *U. humicola* accessions in our previous paper (Higgins et al. 2022) to the assembled chromosomes using Tophat v2.0.13 (Trapnell et al. 2009). The behaviour of this aligner is comparable to Bowtie but designed for RNA reads. A coverage table was generated from the “accepted hits” alignment file using Samtools v1.51.1.

## RESULTS AND DISCUSSION

### Genome assembly

We generated a genome assembly from PacBio HiFi reads alone, using the assembler’s default settings. The resulting primary unitigs (p_utg) had a total length of 4,193 Gbp across 8,785 unitigs. The concatenated contigs from both haplotype files (hap1 + hap2) consisted of 1,087 contigs with a total length of 4,102 Gbp, and a N50 of 32,75 Mbp. 87 % of the genome (3.569 Gb) was assembled into 115 contigs longer than 10 Mbp, and the longest 200 contigs accounted for 97 % (3.98 Gb) of the genome (table 1 and figure 1).

**Figure 1:**
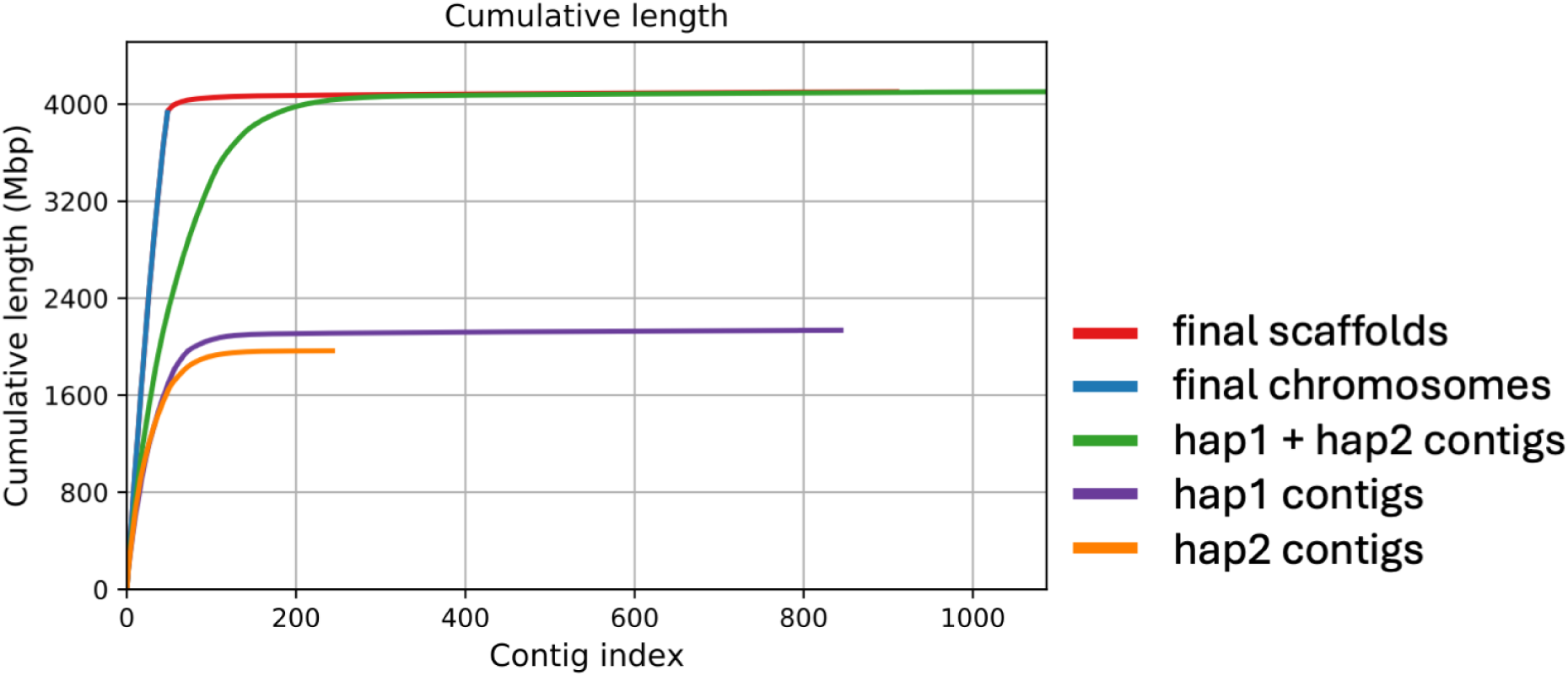
Distribution of cumulative sequence lengths before and after anchoring across different subsets: within the contigs of each haplotype (hap1, hap2), the combined contigs (hap1 + hap2), and in the 48 chromosomes after anchoring, as well as unanchored contigs. Sequences are ordered from longest to shortest along the x-axis.

**Table 1.** Summary statistics of the *Urochloa humidicola* cv. Tully genome at different stages, unitigs, contigs and chromosomes.

| Statistics | Final genome | Chromosomes | Contigs hap1+hap2 | Contigs hap1 | Contigs hap2 | Primary contigs | Unitigs |
| --- | --- | --- | --- | --- | --- | --- | --- |
| Number contigs | 910 | 48 | 1,087 | 844 | 243 | 754 | 8,785 |
| Number contigs (>= 25 kbp) | 888 | 48 | 1,065 | 823 | 242 | 726 | 7,233 |
| Number contigs (>= 50 kbp) | 357 | 48 | 534 | 333 | 201 | 277 | 2,423 |
| Largest contig (Mb) | 106.79 | 106.79 | 98.63 | 98.63 | 95.29 | 93.65 | 56.63 |
| Total length (Gbp) | 4.102 | 3.93 | 4.102 | 2.136 | 1.966 | 2.317 | 4.193 |
| Total length (>= 25 kbp) | 4.101 | 3.93 | 4.101 | 2.135 | 1.966 | 2.317 | 4.16 |
| Total length (>= 50 kbp) | 4.081 | 3.93 | 4.081 | 2.117 | 1.964 | 2.299 | 3.993 |
| N50 | 86.792 | 88.436 | 32.755 | 32.755 | 31.873 | 55.243 | 85.232 |
| N75 | 73.268 | 73.95 | 18.507 | 19.622 | 17.884 | 28.097 | 3.793 |
| L50 | 22 | 21 | 41 | 23 | 19 | 17 | 132 |
| L75 | 35 | 33 | 83 | 44 | 39 | 32 | 314 |
| GC (%) | 45.76 | 45.73 | 45.76 | 45.72 | 45.81 | 45.71 | 45.76 |
| N's | 10,263,851 | 9,114,431 | 10,246,151 | 6,011,896 | 4,234,255 | 0 | 0 |
| N's per 100 kbp | 250.23 | 231.91 | 249.8 | 281.47 | 215.4 | 0 | 0 |

We aligned the Poaceae BUSCO markers to both unitigs and contigs, achieving 99.5% alignment success, with 99.1% being duplicated in both cases. We counted how many times each single-copy BUSCO marker aligned in the contigs (figure 2); 59.2 % of the markers aligned eight times (2899 markers out of 4875), indicating that most haplotypes were present eight times. Assuming a base chromosome number of six, this led to an initial estimation of 48 chromosomes.

**Figure 2:**
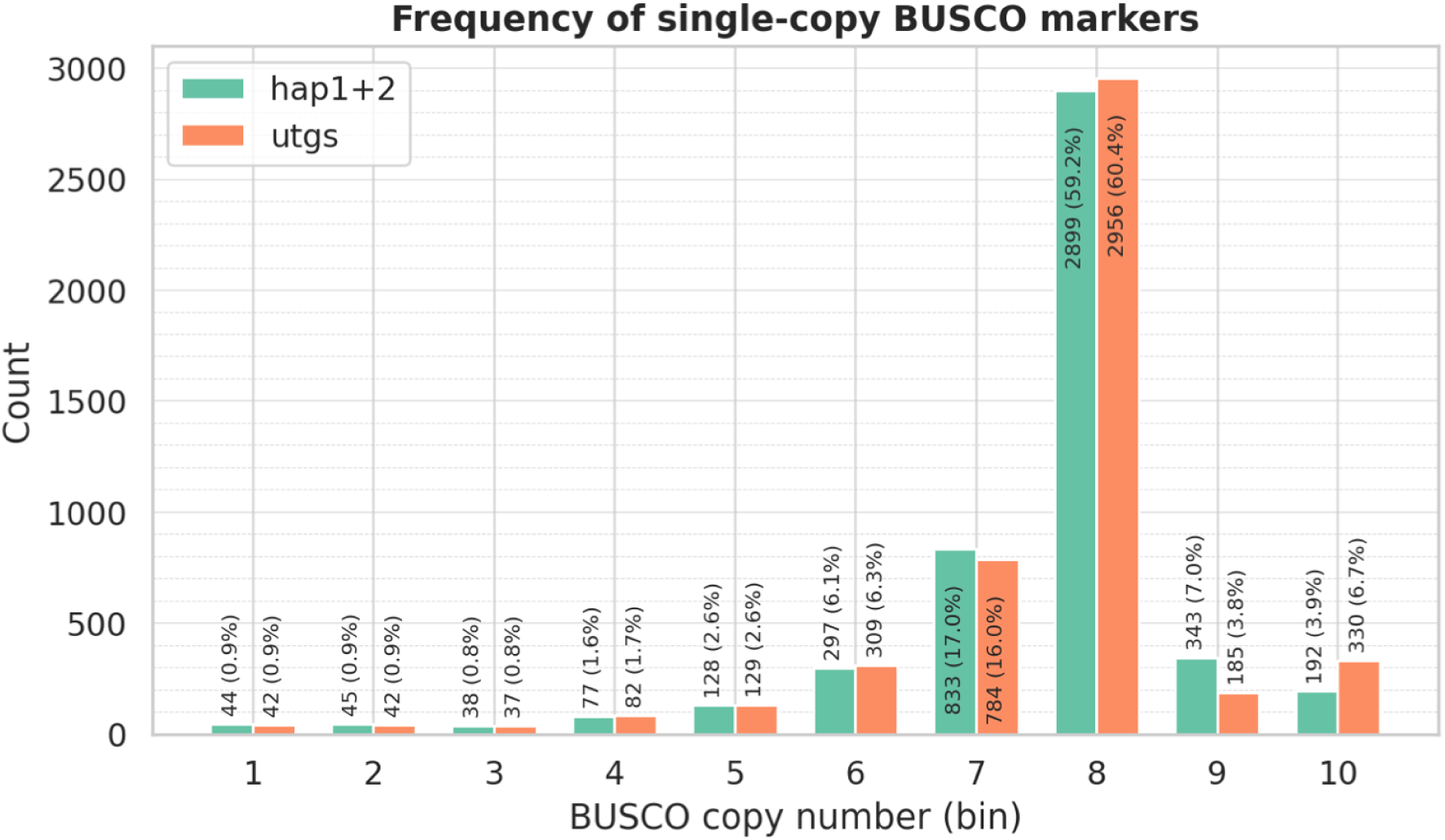
Distribution of BUSCO copy numbers across unitigs (utgs) and contigs (hap1+hap2). The histogram shows the number of alignments for each BUSCO marker, indicating that about 59% of the markers are present eight times, consistent with an octoploid genome structure.

Since 97% of the genome was assembled into 200 contigs (figure 1), and we expected 48 chromosomes, we concluded that most chromosomes were assembled in just a few contigs (around four on average). Consequently, we decided to scaffold them using synteny, as detailed in the methods; firstly, comparing the hap1 and hap2 contigs to the primary contigs assembled by Hifiasm, then by comparing the “hap1” to the “hap2” homologs to each other, and finally by comparing to the longest assembled haplotype in each chromosome (often a full chromosome at this stage). In total, 225 contigs were scaffolded in 48 chromosomes. Three chromosomes were assembled end-to-end in one contig, and nine more in two contigs. Only six chromosomes were assembled in seven or more contigs.

We annotated 259,254 genes and 277,99 transcripts. 249,665 genes (96.3%) were in chromosomes, and the rest were in contigs.

### Differences between ancestries

Chromosomes were assigned to ancestries based on differential k-mer analysis among chromosome sets. The SubPhaser tool clusters the subgenomes using k-means, from which it constructs a hierarchical dendrogram and heatmap (figure 3A) based on the presence-absence variation of these differential k-mers, as well as a PCA (figure 3B). The composition of cv. Tully was AABBBBCC for chromosomes 1 to 4, and AAABBBCC for chromosomes 5 and 6. This can also be expressed as 14A 22B 12C, or 12A+2 24B-2 12C.

**Figure 3:**
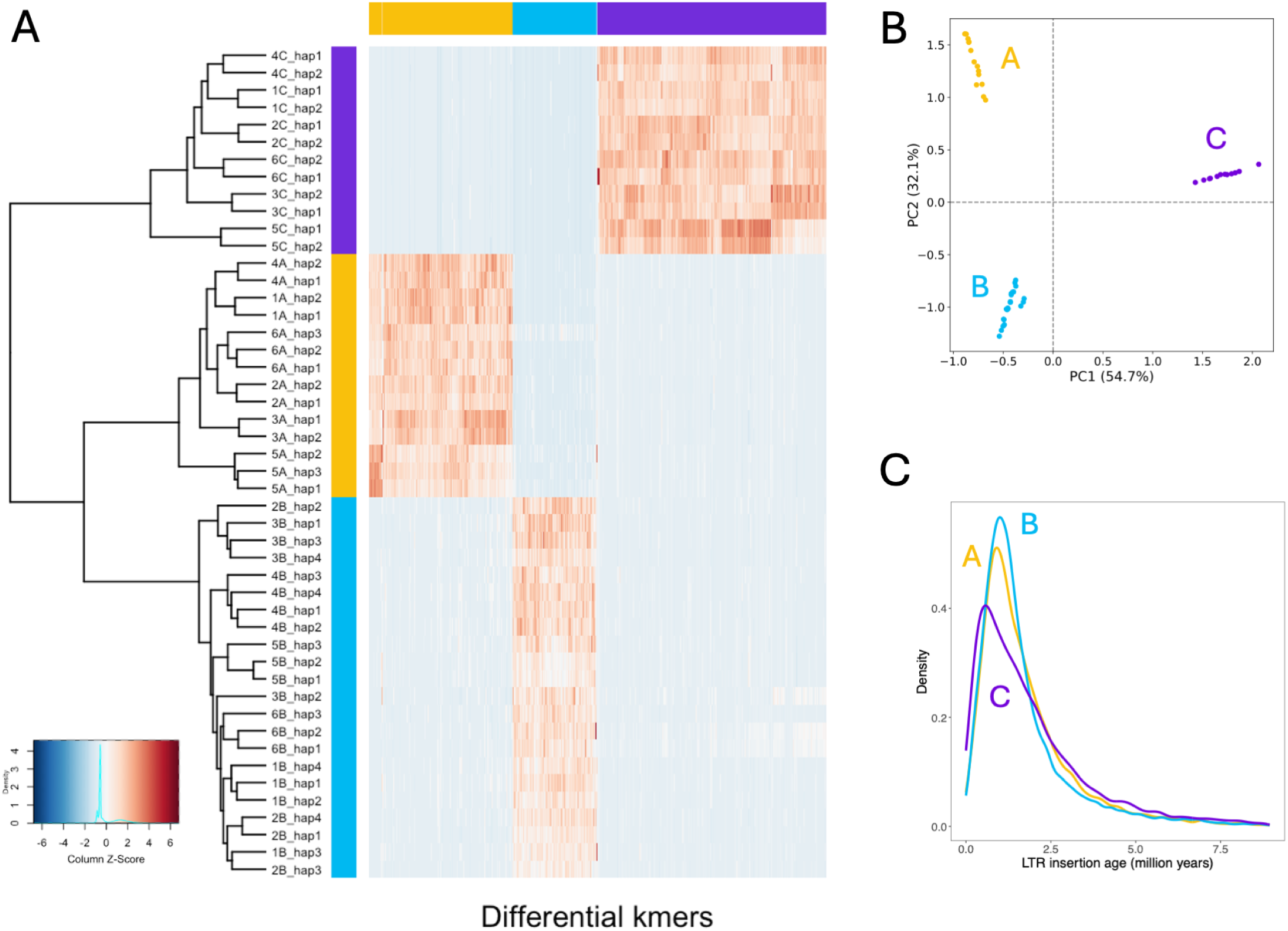
Subgenome classification of the 48 chromosomes in *U. humidicola*. (A) Hierarchical clustering heatmap based on differential Kmers, and (B) Principal-component analysis (PCA) based on differential Kmers. Both analyses differentiate chromosomes into three subgenomes (A, B, and C). (C) Insertion-age distribution of LTR-retrotransposons reveals overlapping peaks for A and B and older insertions for C, pointing to distinct evolutionary histories.

A haplotype analysis based on chromosome alignment showed the shared synteny between subgenomes (figure 4). The A and B copies of each chromosome were highly similar (green links in figure 4). The C subgenome exhibited lower synteny (red links in figure 4). In all chromosomes, there was no synteny with the centromeric regions on the C subgenome chromosomes; this is particularly clear in chromosomes 3, 4 and 5. Synteny between C and the other two subgenomes was particularly low in chromosome 5.

**Figure 4:**
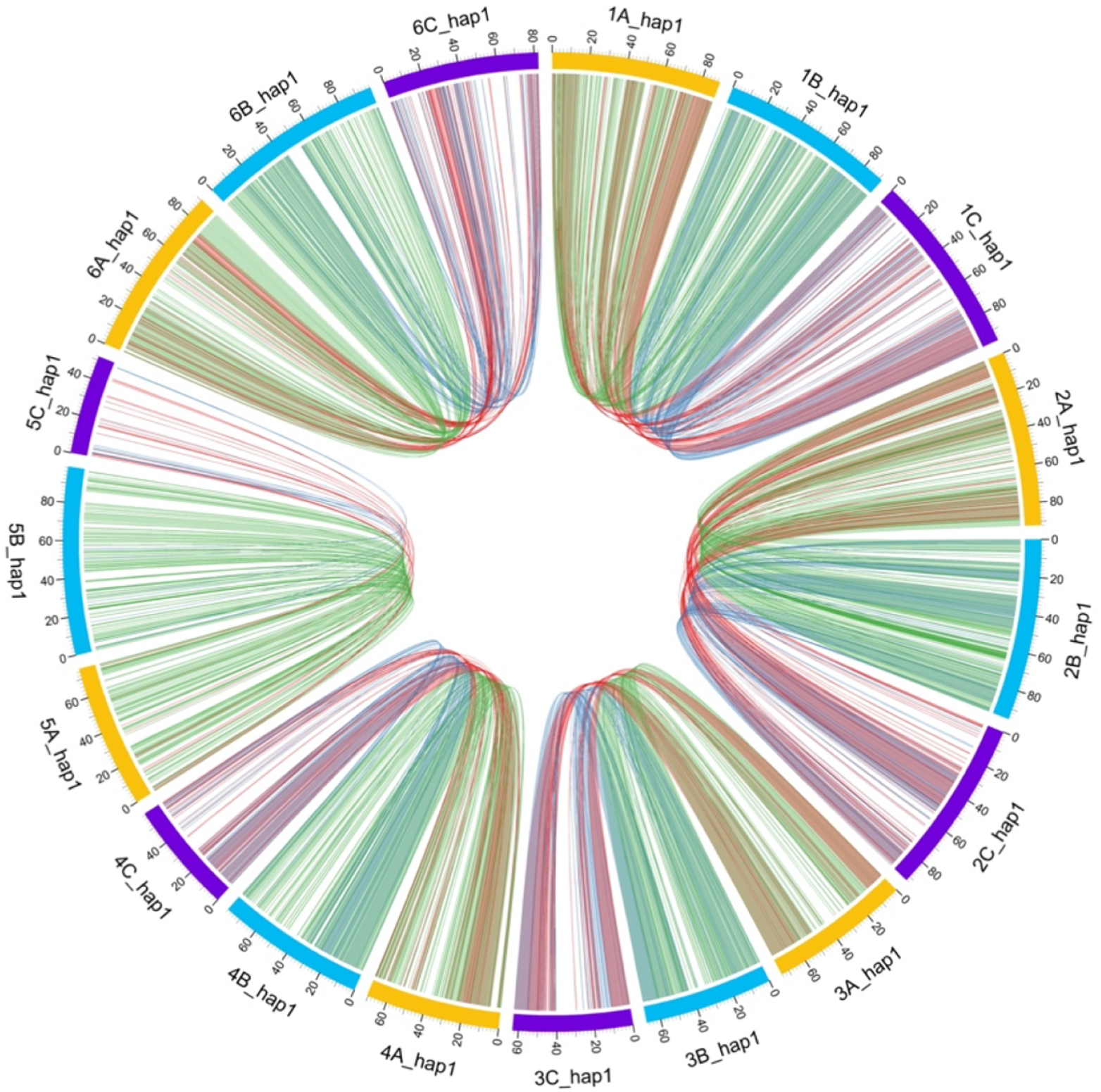
Comparative synteny among subgenomes. Circos plot illustrating shared syntenic blocks among the A, B, and C subgenomes. Green ribbons represent A–B synteny, yellow ribbons denote B–C, and red ribbons show A–C relationships; reduced collinearity of C-subgenome chromosomes is evident near centromeres.

Subphaser was also used to analyse the LTR-RT elements by subgenome by enrichment analysis of LTR-RTs detected by several classification tools. The insertion age was estimated per subgenome (Figure 3C), which supported a closer relation between subgenomes A and B than with C. We also obtained phylogenetic trees of subgenome-specific LTR/Gypsy and LTR/Copia elements (figure 5). In most LTR clades, there was a clear pattern where A and B clustered together, sometimes mixed, and C clustered separately.

**Figure 5:**
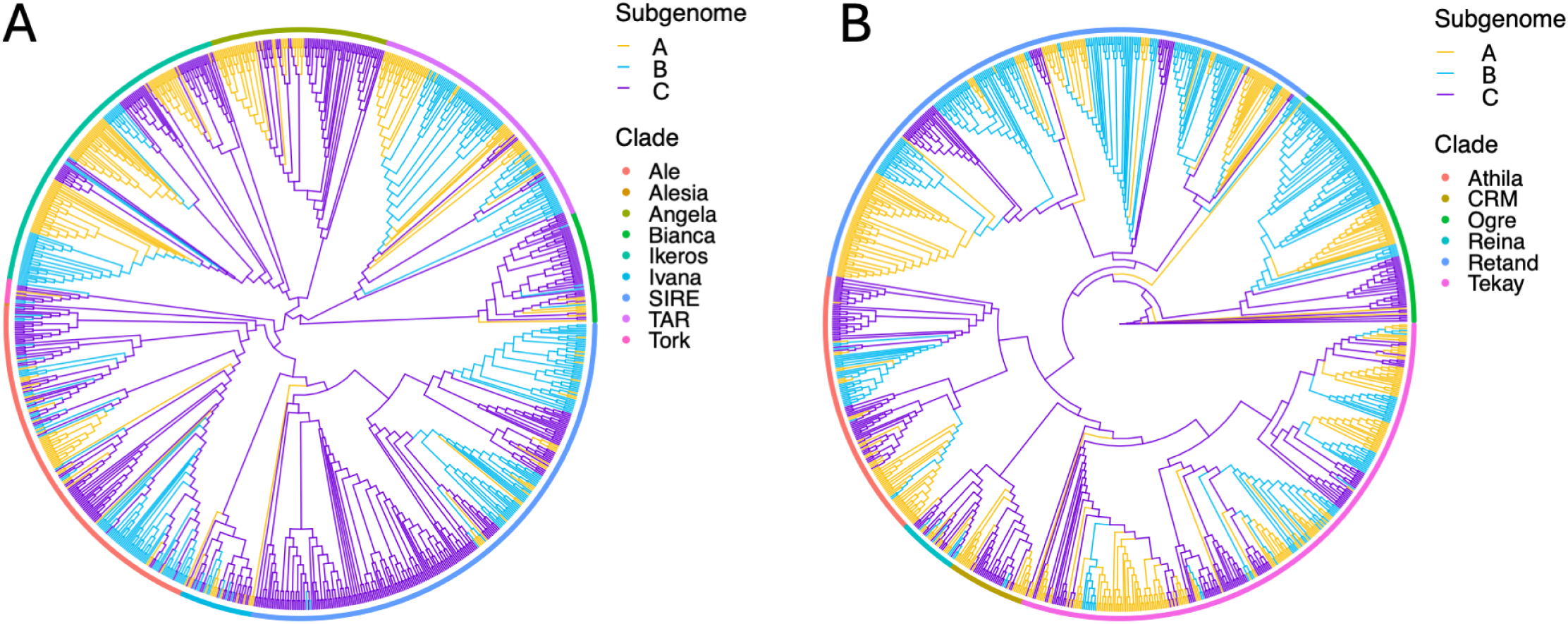
Distribution and phylogeny of LTR-retrotransposons. Bar plots show the relative abundance of Gypsy and Copia elements in each subgenome, and maximum-likelihood trees depict their relationships. Clustering of A and B LTR lineages and separation of C elements support a shared transpositional history between A and B subgenomes.

The clear clustering of A and B subgenomes in both differential k-mer and LTR analyses indicates that these two ancestries diverged more recently from each other than from the C genome donor. The shared peaks of amplification for Copia and Gypsy elements suggest that A and B lineages experienced similar transpositional bursts, possibly during a common polyploidization event. In contrast, the C subgenome shows a different repeat composition and older insertion ages, which align with a long history of independent evolution before its introgression. Collectively, these findings support a stepwise allopolyploid origin for *U. humidicola*, with the A–B component forming first, followed by the later addition of the diverging C genome.

### Ancestral origins

Most *Urochloa* species, particularly wild ones, lack WGS data. This led us to use the “Angiosperms 353” (A353) markers from the “Tree of Life” project, which had been generated for at least 15 *Urochloa* species (Baker et al. 2022; Masters et al. 2024). For this, we identified and extracted the homologs to the A353 marker from the new genome in each *U. humidicola* subgenome and haplotype. The tree we obtained (figure 6) consistently placed the B subgenome with *U. dictyoneura* and separated it from other CWRs in the same clade. This suggests that *U. dictyoneura* is the ancestor of the B subgenome. These two species are known to intercross and share the same base chromosome number of six. The A subgenome was included in the Humidicola clade, but was not one of the wild relatives for which we had markers. The C subgenome is phylogenetically distant from the previous, and is likely *U. arrecta* from the Mutica clade (figure 6).

**Figure 6:**
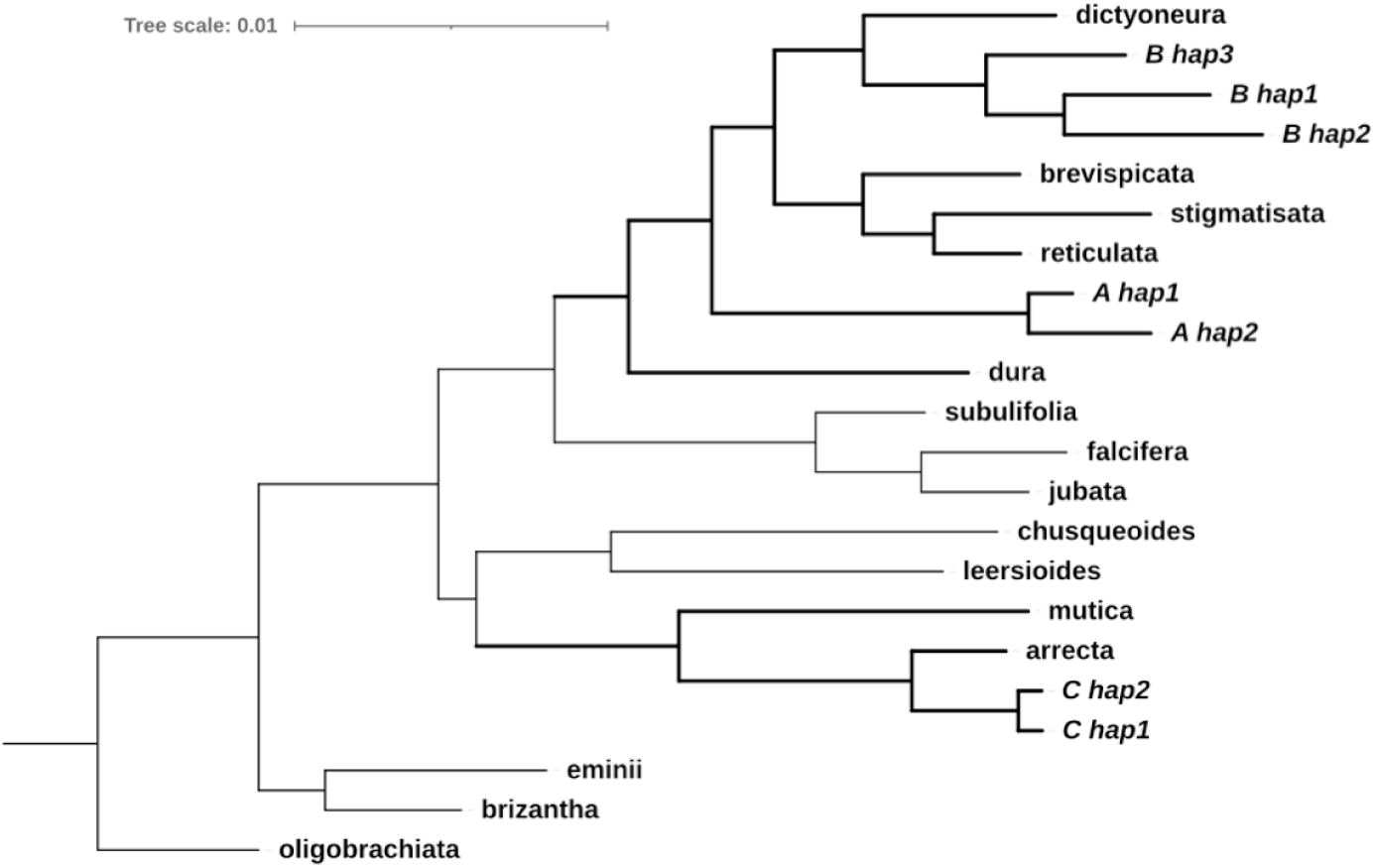
Phylogenetic placement of *U. humidicola* subgenomes among related *Urochloa* species. The maximum-likelihood tree, constructed using 353 protein markers (“Angiosperms 353”), shows the B subgenome grouping with *U. dictyoneura* and the C subgenome with *U. arrecta*, while A subgenome sequences cluster independently within the Humidicola clade.

This phylogenetic framework implies a historical hybridization across clade boundaries, underscoring the reticulate nature of *Urochloa* evolution. These relationships provide a robust basis for revising taxonomic relationships within the species and genus. Our results highlight an evolutionary trajectory of repeated hybridisation events, at least as complex as in the Brizantha complex.

### Genomic composition in *U. humidicola*

We aligned the publicly available DNA-seq and RNA-seq data from *U. humidicola* to the reference genome. Our alignment parameters allowed each read to align only to its “top best” position (figure 7). CIAT-26155 (SRR16327314) underwent whole-genome resequencing and was identified as hexaploid through k-mer analysis (Tomaszewska et al. 2023). Among the reads, 15,646,090 aligned to a chromosome from the A subgenome (33,32%), 31,076,742 to a chromosome from the B subgenome (66,19%), and 230,974 to a chromosome from the C subgenome (0,005%). The ratio of reads between the A and B subgenomes was clearly 1A:2B, supporting a composition of 12A and 24B (AABBBB) for CIAT-26155. Conversely, accession 16888, expected to be an aneuhexaploid (Worthington et al. 2019), showed about 40% of the reads aligned to A subgenome chromosomes and roughly 60% to B subgenome chromosomes, resulting in a 1A:1.5B ratio. This indicates an overrepresentation of A chromosomes compared to B, but could also be explained by unbalanced marker density, as this accession was genotyped using Genotyping-By-Sequencing (GBS). An 1A:1.5B ration would be equivalent to the 14A 22B composition observed in the genome of cultivar Tully.

**Figure 7:**
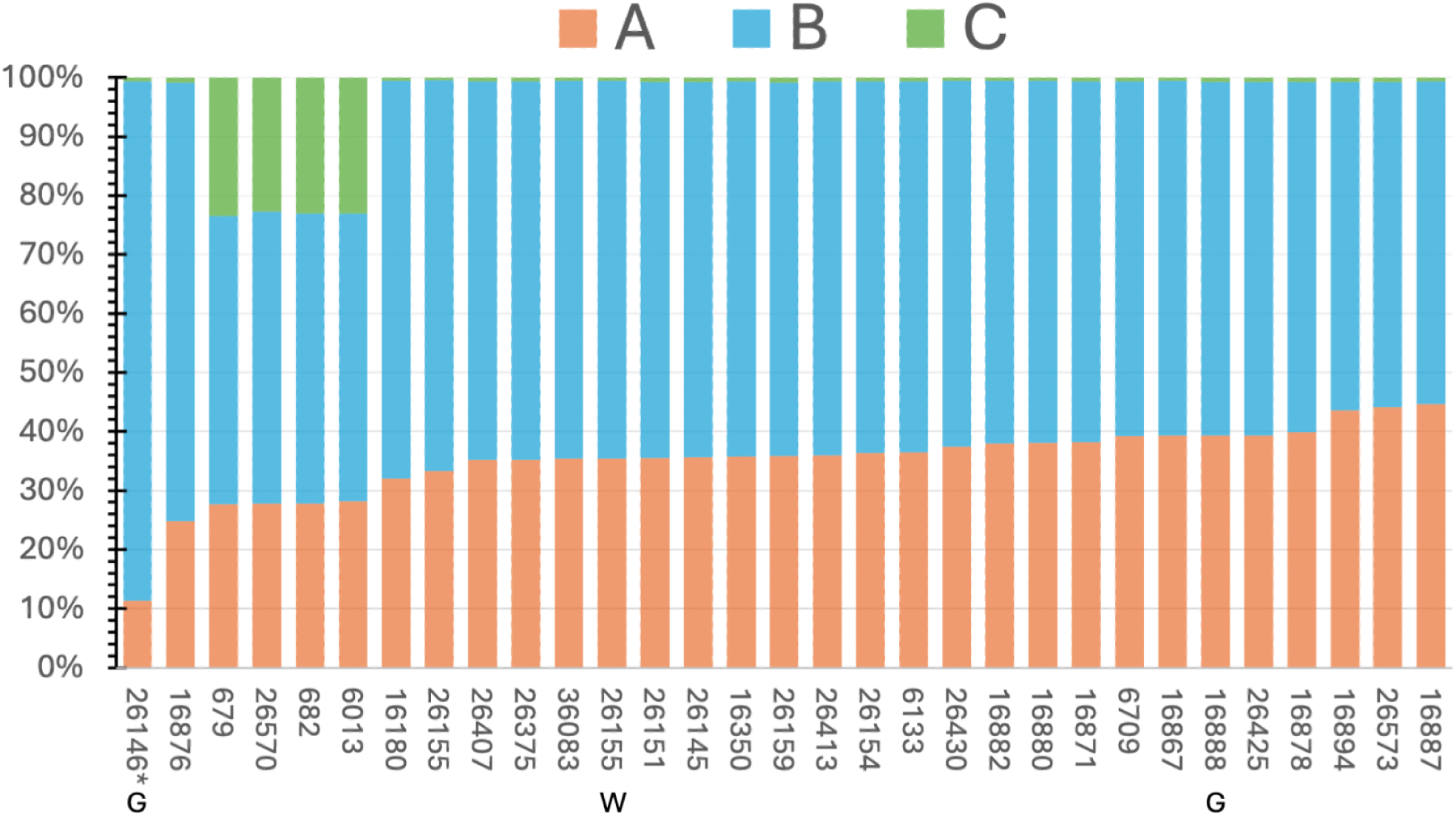
Competitive read mapping of public *U. humidicola* accessions to subgenomes. Stacked bar plots show the proportion of reads aligning to chromosomes of the A, B, and C subgenomes for each accession. They were genotyped using RNA-seq except 26155 (labelled W for WGS), and 26146 and 16888 (labelled G for GBS). These ratios mainly reflect the variability in subgenome copy numbers and ploidy levels across different accessions.

The competitive mapping results for the sexual accession CIAT-26146 (Worthington et al. 2019) were markedly different, with only 10% of reads aligning to the A subgenome and 90% to the B subgenome (figure 7), resulting in a 1:9 ratio between A and B. This accession was also genotyped using Genotyping-By-Sequencing (GBS), an as before, there may be unbalanced marker density. Accepting the likely hexaploidy (Worthington et al. 2019), this supports an autoploid background (BBBBBB), ranging from none to up to three A chromosomes (0-3A, 36-33B).

RNA-seq reads (Higgins et al. 2022) from four accessions (figure 7), including cv. Tully itself (labelled as CIAT-679), aligned to the C subgenome with a frequency of around 23%. None of the remaining accessions aligned to the C subgenome (<0.01% of the reads). These four accessions aligned to the A and B subgenomes with similar proportions, so we should assume a composition similar to cv. Tully. In Higgins et al. 2022, these four accessions were grouped into a different subpopulation than the other accession, called “humidicola-2” (Higgins et al. 2022). This evidence supports that these subpopulations were split by the presence or absence of C ancestry.

Among the remaining accessions, most displayed a ratio close to 1A:2B (about 33% reads aligned to A and 66% reads aligned to B), indicating that the most common composition is AABBBB. However, their ratios ranged from 1A:1.25B in CIAT-16887 to 1A:3B in CIAT-16876 (figure 7). While coverage analysis from RNAseq is noisier than WGS, as it reflects allelic expression ratios rather than homolog rations; this variation could equally indicate differing ploidy levels beyond hexaploidy; for example, a composition of AABBBBBB would match the ratio observed in CIAT-16876.

The diversity of subgenomic ratios among accessions indicates ongoing genomic fluidity in *U. humidicola* populations, reflecting recurrent hybridization and uneven chromosome transmission. Such variation may underlie differences in reproductive mode, as apomictic accessions maintain fixed heterozygosity, whereas sexual forms may re-shuffle subgenomic composition through residual meiotic pairing.

### Evolution of *U. humidicola*

We propose a model (figure 8) that summarises the inferred origin and diversification of *U. humidicola* cytotypes and accessions based on the analysis of this genome reference and competitive read-mapping against it. The ancestors *U. dictyoneura* (2n = 4x = 24; B genome), wild *Urochloa* from the Humidicola clade (A genome), and *U. arrecta* (2n = 4x = 36; C genome) are shown at the top. U. dictyoneura is likely autopolyploid (reference), but it could also be allotetraploid.

**Figure 8.**
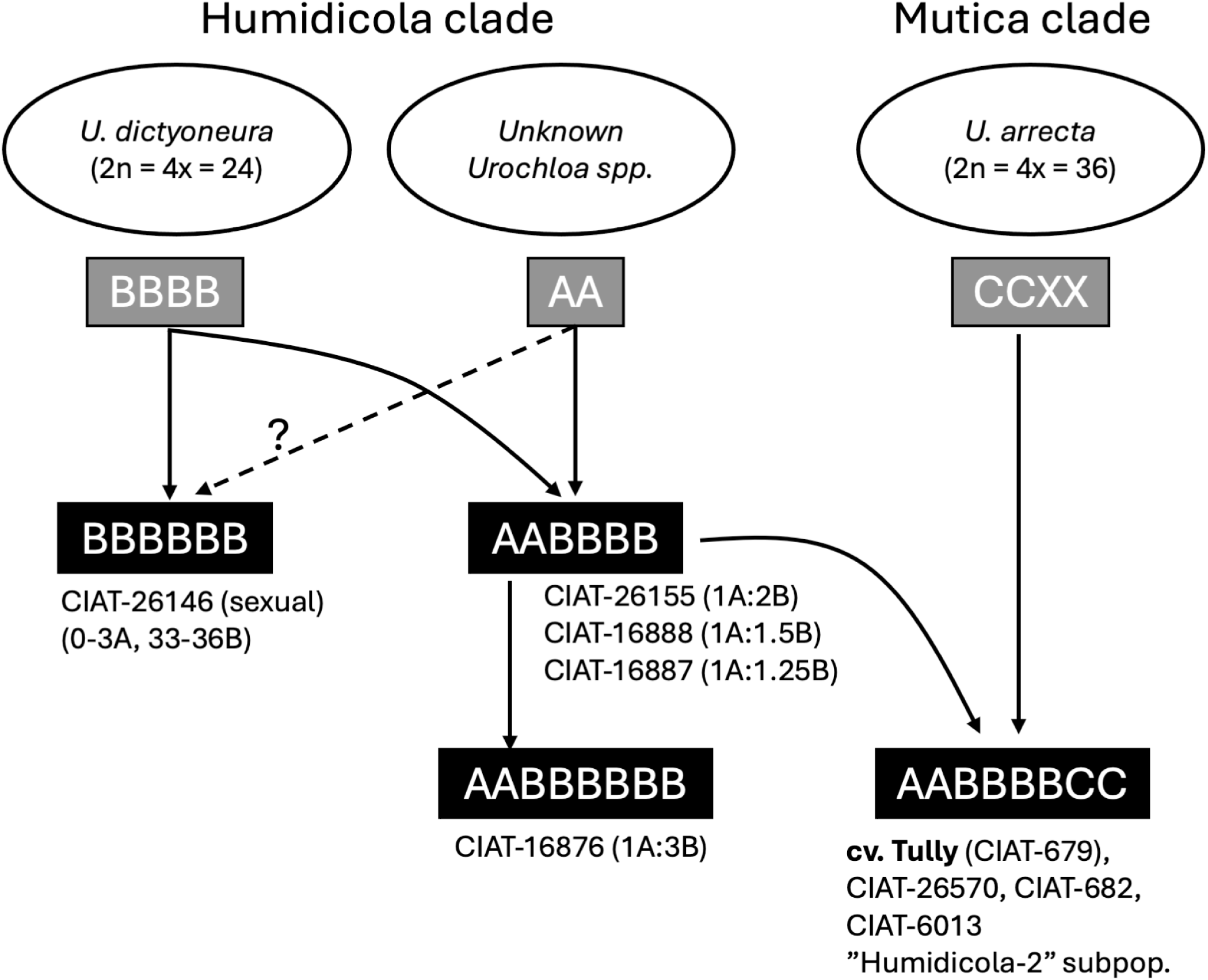
Evolutionary model of the composition among *Urochloa humidicola* accessions and putative events leading to different *U. humidicola* cytotypes (black boxes), reflecting the species’ complex reticulated evolution. The B-genome donor *U. dictyoneura*, an A-genome wild *Urochloa* from the Humidicola clade contributed to the common hexaploid (AABBBB), as well as increasing ploidies, such as octoploid (AABBBBBB). While *U. arrecta* (C genome) contributed to AABBBBCC lineages, including the apomictic cultivar Tully sequenced.

Initial hybridisation between *U. dictyoneura* (or a very closely related species) and the unknown A genome donor produced the AABBBB-type hexaploids, which are the most common types in the genebank collection. Selected examples include accessions CIAT 26155, CIAT 16888, and CIAT 16887, and variation in the ratios obtained can reflect additional copies of A-origin chromosomes, as observed in cv. Tully. A second lineage derived from *U. dictyoneura* alone generated BBBBBB cytotypes, such as the sexual accession CIAT-26146. Further hybridisation among AABBBB cytotypes likely produced octoploid forms (AABBBBBB), exemplified by CIAT-1673 (12A 36B).

A subsequent introgression event involving the C genome from *U. arrecta* (or a very closely related species) led to the creation of the cultivar *U. humidicola* cv. Tully (AABBBBCC), which serves as the archetype of the “humidicola-2" subpopulation. This subpopulation also includes CIAT-26570, CIAT 682, and CIAT 6013. These accessions retain genetic signatures of the A, B, and C subgenomes, aligning with the structure observed in this cv. Tully assembly.

The model demonstrates how recurrent hybridisation, partial diploidisation, and subgenome retention have shaped the mosaic ancestry of *U. humidicola* and explain the genomic diversity observed among apomictic and sexual accessions. The inferred hybridisation network (Figure 8) reflects the broader pattern of reticulate evolution observed across tropical forage grasses, where polyploidy and apomixis repeatedly facilitate the stabilisation of interspecific hybrids. Such processes parallel those seen in *Paspalum* and *Poa*, suggesting convergent evolutionary solutions to reproductive assurance under tropical conditions.

## Conclusion

The identification of distinct A, B, and C ancestries in *U. humidicola* now enables subgenome-specific analyses of gene expression, methylation, and apomixis regulation, which were previously obscured by the lack of a complete reference. Researchers working with hexaploid *U. humidicola* panels or families simply need to remove the C ancestry chromosomes to obtain an accurate reference for their system. The haplotype-complete assembly offers a framework to locate and characterise the chromosomal regions associated with apomixis, previously mapped only through low-resolution markers (Worthington et al. 2019). Beyond evolutionary insights, the genome provides practical targets for breeding. The clear delineation of subgenomes allows genome-wide association studies and marker development for traits such as forage digestibility and biological nitrification inhibition (BNI). These advances align with global efforts to enhance tropical pasture productivity while reducing methane emissions and nitrogen losses, positioning *U. humidicola* as a model for genomic strategies toward low-carbon livestock systems. This genome represents a foundational resource for the *Urochloa* research community. It supports future comparative genomics across tropical forage species, informs efforts to engineer apomixis in crops, and contributes to global initiatives for climate-resilient agriculture.

## Data Availability Statement

All data are deposited in the SRA under accession PRJEB90424. The genome assembly, together with its gene annotation, was deposited in ENA with accession GCA_965614515.2 (https://www.ebi.ac.uk/ena/browser/view/GCA_965614515.2). The scripts used are available at https://github.com/DeVegaGroup/Assembly-and-analysis-Urochloa-humidicola-genome.

## Acknowledgments

All the authors approved this manuscript. We thank the Australian Pastures Genebank and the CIAT CGIAR Genebank platform in Colombia for providing the accessions. The authors would like to acknowledge the support of the Transformative Genomics group at the Earlham Institute, especially Drs. Sacha Lucchini, Leah Catchpole, Karim Gharbi, and Chris Watkins for their comprehensive support and roles in platform management and supervision. We also acknowledge the invaluable support from the Earlham Institute’s research office, BDE and support teams, the Norwich Bioscience Institute’s research computing and contracts teams, and horticultural services at the John Innes Centre.

